# T cell-microbiome associations captured through T cell receptor convergence analysis

**DOI:** 10.1101/2025.09.30.679432

**Authors:** Romi Vandoren, My Ha, Vincent Van Deuren, Naomi De Roeck, Ting Pu, Eelco C. Brand, Maria Kuznetsova, Hajar Besbassi, Esther Bartholomeus, Fabio Affaticati, Ilke De Boeck, Thies Gehrmann, Sarah Lebeer, Bas Oldenburg, Femke van Wijk, Peter Delputte, Sara Verbandt, Sabine Tejpar, Kris Laukens, Benson Ogunjimi, Pieter Meysman

**Author notes:** **Corresponding author** (R.V.) and (P.M.). Shared senior authors (Benson Ogunjimi, Pieter Meysman).

## Abstract

The gut microbiome modulates mucosal immunity, yet how specific bacterial taxa shape the diversity and specificity of T cell receptor (TCR) repertoires remains poorly understood. Existing approaches emphasize single-species effects or broad immune features, without pinpointing which microbes drive specific T cell clonotypes. We present AIRRWAS, a computational framework that integrates TCR-microbiome interaction analysis with targeted *in vitro* validation to detect genus-level TCR convergence. Applied to three independent cohorts, AIRRWAS identified reproducible associations between convergent TCR clusters and 21 bacterial genera spanning core commensals, probiotics and taxa with immunomodulatory roles. Predicted clonotypes were enriched within the TCR–microbiome interaction network and preferentially activated by genus-matched stimuli, eliciting different functional T cell responses. These findings demonstrate that distinct repertoires can share genus-specific TCR motifs, enabling detection of shared immune signatures. AIRRWAS can map these TCR– microbiome interactions, laying the groundwork for biomarker discovery immune monitoring and the development of microbiome-targeted therapies.

## Introduction

Mucosal immunity is essential for preventing systemic infections, chronic inflammation and maintaining immune homeostasis^1^ in the gastrointestinal^2–4^, respiratory^5^, and urogenital^6^ tracts. Within this system, T cells play a central role in responding to the microbial environment. Dysregulation of these responses contributes to inflammatory disorders such as inflammatory bowel disease (IBD) and colorectal cancer (CRC)^7–9^. Conventional effector T cells, including CD4⁺ T helper subsets (Th1, Th17) and CD8⁺ cytotoxic T cells, mount protective responses against invading pathogens, while regulatory T cells (Tregs) suppress excessive inflammation to maintain immune tolerance toward commensal species^9, 10^. T cell differentiation, activation and function is strongly influenced by the gut microbiome, highlighting the need to understand how T cells respond to microbial antigens^9, 11–13^. At the core of this interaction is the T cell receptor (TCR), a heterodimer of α and β chains, that determines antigen specificity and mediates recognition of microbial peptides. Previous research has shown that microbiome-specific TCRs can be found both in local mucosal tissue and peripheral blood, although more abundant in the tissue^14–16^.

Despite their importance, TCR-microbiome interactions have been difficult to characterize. High-throughput TCR sequencing has revealed the enormous diversity of TCR repertoires but linking them to microbial antigens is complicated by inter- and intra-individual variation in both microbiome composition and TCR repertoire^17, 18^. Previous approaches mainly rely on community-level or single-species analysis in combination with high-level TCR metrics like repertoire diversity or clonality^19–27^. While interesting, these methods capture only broad correlations but miss clonotype-specific responses that directly reflect microbial recognition. Single-cell and antigen screening methods have provided important insights but their scalability and reproducibility across large cohorts remain limited. As a result, fundamental questions regarding which bacterial taxa shape TCR clonotype signatures and whether these associations can reveal immunomodulatory microbes or disease-linked responses remain.

To overcome these challenges, we developed Adaptive Immune Receptor Repertoire-Wide Association Study (AIRRWAS), a computational framework that integrates microbiome data (from stool or tissue) with high-throughput TCR sequencing (from blood or tissue) to systemically link bacterial taxa to TCR clonotypes. AIRRWAS identifies convergent TCRs associated with bacterial genera across individuals, sequencing platforms and data types. By operating not only at the clonotype but also the cluster level, it overcomes the limitations of repertoire-level analyses and allows for detection of microbial drivers of T cell activation and tolerance that are invisible to conventional approaches.

We show that AIRRWAS enables systemic, high-resolution mapping of TCR-microbiome interactions. The framework provides insight into how the microbiome shapes immunity and identifies reproducible, convergent TCR clonotypes consistently linked to specific bacterial genera. By integrating additional data modalities, we can associate clonotype-specific responses with mechanistic insights, providing a functional level of interpretation. Together, these results demonstrate that AIRRWAS uncovers robust, biologically meaningful TCR-microbiome associations and provides a flexible strategy for exploring microbiome-immune dynamics. Beyond the gut, the framework can also be applied to other mucosal surfaces, offering a generalizable approach for linking microbial composition to T cell immune responses and informing pathophysiological insights, biomarker discovery, patient stratification and the development of targeted interventions to modulate immune responses in health and disease.

## Results

### TCR diversity, epitope prediction and microbiome profiling

We analyzed the TCR repertoires in a discovery cohort of 74 individuals with PBMC TCR sequencing and 16S rRNA microbiome data (Extended Data Fig. 1). The dataset comprised >6 million TCR sequences, including unpaired α chains (3,107,792) and β chains (2,974,267) from CD4^+^ (4,911,201) and CD8^+^ (1,170,858) T cells. Diversity metrics (richness, Pielou’s evenness, Gini-Simpson index, Gini coefficient and DE25) were compared between different groups of TCRs (TRA-CD4, TRB-CD4, TRA-CD8 and TRB-CD8), showing a trend towards lower richness and diversity for CD8^+^ cells compared to CD4^+^ cells but higher levels of expansion (Figure 1a). Beta diversity analysis using Bray-Curtis dissimilarity revealed no grouping of samples at the repertoire level (Extended Data Fig. 2a). Epitope specificity predictions (ImmuneWatch Detect, v1.4.0) identified 153,763 TCRs (2,5%) predicted (score ≥ 0.2) to recognize 537 epitopes from 184 distinct antigens, classified as viral, human, microbial or synthetic. The majority targeted Influenza A virus, SARS-CoV-2 or human antigens, followed by HIV, CMV and EBV (Extended Data Fig. 2b). A total of 6,594 TCRs were predicted to recognize bacterial epitopes, primarily α chains with mucosa-associated invariant T (MAIT) cell-like V- and J-gene usage, targeting MR1:5-OP-RU (Riboflavin, Vitamin B2, n=5,915), ECTGLAWEWWRTV (Pneumolysin, *Streptococcus pneumoniae*, n=483), VMATRRNVL (p55, *Mycobacterium tuberculosis*, n=121), VMTTVLATL (p34, *Mycobacterium tuberculosis*, n=59) and PMPMPELPYP (PFSGDS, *Pseudomonas aeruginosa*, n=16).

**Figure 1.**
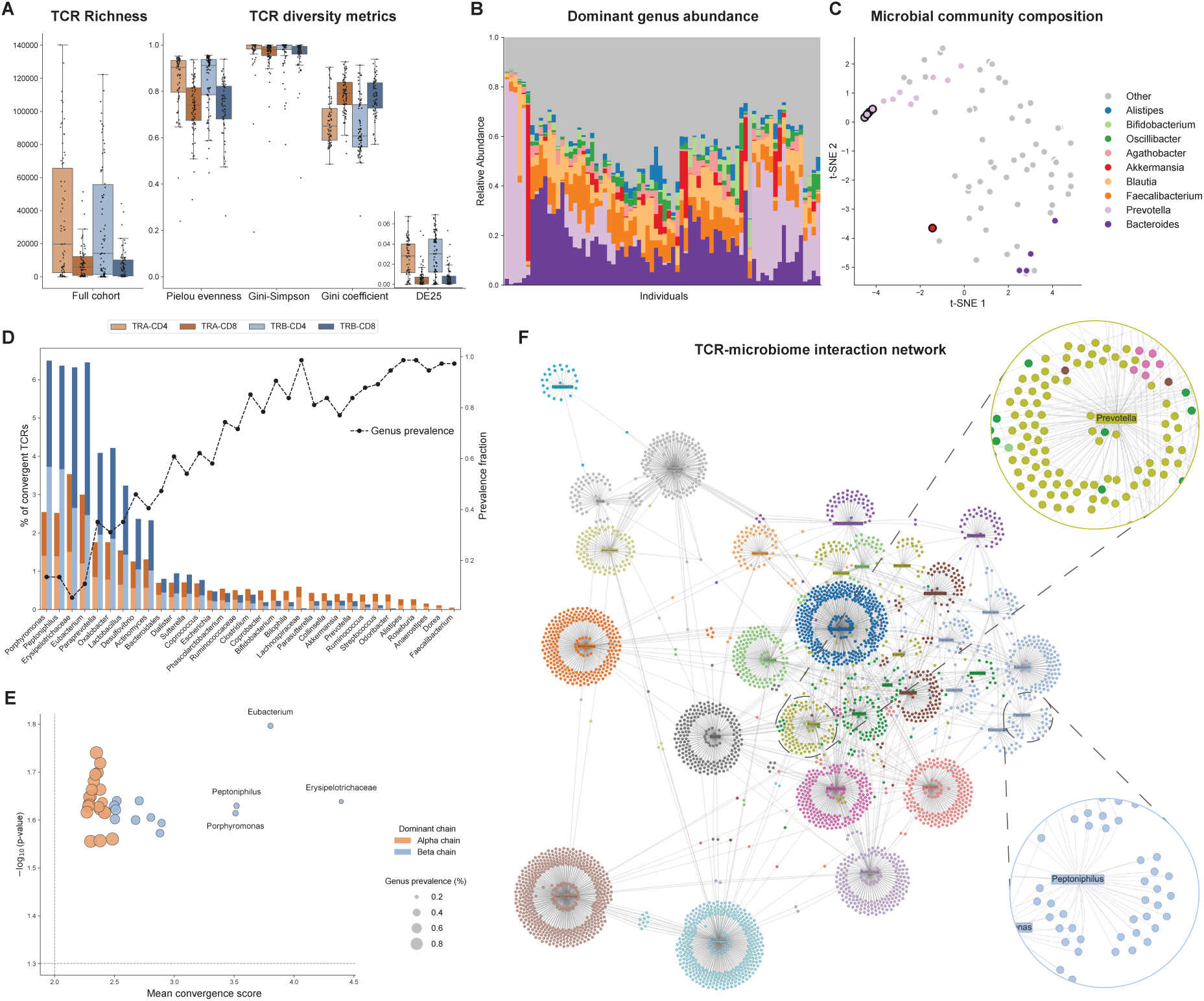
**AIRRWAS reveals genus-specific TCR clonotypes and motifs**. **a**, TCR repertoire analysis across chains and subgroups. Left boxplots show repertoire richness for TRA-CD4, TRA-CD8, TRB-CD4, and TRB-CD8 across the full cohort. Right boxplots display several repertoire diversity metrics (Pielou’s evenness, Gini-Simpson diversity, Gini coefficient and DE25) for the same subgroups. There are no differences between α and β chains, but we do observe an increased repertoire diversity for the CD4^+^ T cell subgroup, while CD8^+^ showed higher clonal expansion measured by DE25. **b**, Relative abundance of the top nine microbial genera per sample, with remaining genera aggregated as “Other.” Each bar represents an individual sample. **c**, t-SNE projection of microbial beta diversity profiles colored by the most abundant genus. Grey dots indicate samples dominated by multiple low-abundance genera. Black outlines indicate that the top genus comprises >50% of the total abundance in the sample. The even distribution suggests no major clustering driven by genus-level dominance. **d**, Stacked barplot of convergent TCR fractions stratified by chain (TRA/TRB) and subset (CD4/CD8), with a line showing genus prevalence across samples. **e,** Volcano plot showing mean TCR convergence score versus enrichment p value for the identified convergent TCRs for each genus. Dot color indicates the dominant TCR chain (α and β), and dot size reflects prevalence across the discovery cohort (% of patients). Less prevalent genera show β chain dominance and higher convergence. **f**, Network of significant convergent TCR-genus interactions. Nodes represent public TCRs (present in ≥5 individuals) or microbial genera, colored by community determined using Louvain community detection. Edges denote statistically significant convergence between TCR and genus nodes. Two genus clusters (*Prevotella*, *Peptoniphilus*) are highlighted from the network. TRA = TCR α chain, TRB = TCR β chain, DE = diversity evenness

Paired 16S rRNA sequencing detected 3,803 unique ASVs, of which 1,132 were shared between at least 2 individuals. Genus-level profiles revealed large inter-individual variability, five samples were dominated (genus abundance >50%) by a single genus (four by *Prevotella* and one by *Akkermansia*), whereas others showed more balanced communities (Figure 1b). Single-genus dominance, particularly by *Prevotella*, has been reported previously^28^, although its biological basis and implications remain unclear. Beta diversity analysis with Bray-Curtis dissimilarity showed broad sample distribution across the space. Although some single-genus-dominated samples clustered together, most had diverse communities that were more dispersed with no sample stratification (Figure 1c).

### TCR convergence patterns were found associated to bacterial genera

To identify associations between convergent TCRs and bacterial genera, we applied the AIRRWAS framework to paired TCR-microbiome data from all 74 individuals in the discovery cohort (Extended Data Fig. 1). Across 36 genera tested, 33 were associated with significantly convergent TCRs (convergence > 2, p ≤ 0.05). The number of convergent TCRs varied widely between genera and correlated with genus prevalence (Figure 1d). Low-prevalence genera, including *Porphyromonas*, *Peptoniphilus*, *Erysipelotrichaceae* and *Eubacterium*, each had >100,000 convergent TCR hits, whereas high-prevalence genera (>95%) yielded fewer associations (Extended Data Table 1), likely reflecting reduced statistical power when “negative” samples are limited. Mean convergence and enrichment per genus showed that convergent TCRs for low-prevalence genera were dominated by β chains and had a higher convergence score in comparison to high-prevalence genera that are mainly dominated by convergent α chains with lower convergence (Figure 1e).

Publicity analysis, defined as the number of individuals sharing each clonotype, revealed that most convergent TCRs were private, while a subset was shared across multiple individuals with varying level of convergence. Convergence analyses, stratified by chain type and T cell subtype, showed comparable mean convergence between α and β chains and between CD4^+^ and CD8^+^ repertoires. However, increased mean convergence was observed in the low-prevalent genera, while decreased convergence and increased publicity was found linked to high-prevalence genera. This could indicate that some convergent TCRs linked to low-prevalent genera may arise from high generation probabilities rather than genus-specific responses (Extended Data Fig. 2c).

Next, we used AIRRWAS to generate a TCR-bacterial genus interaction network restricted to convergent clonotypes shared by ≥5 individuals, comprising 4,055 significant associations between 3,535 unique TCRs and 33 genera (Extended Data Table 2). Nodes represent either convergent TCRs or bacterial genera, and edges indicate significant associations (Figure 1f). The network contained highly connected hubs, including *Lachnospiraceae* (family), *Erysipelotrichaceae* (family) and *Odoribacter*, which were linked to the greatest number of public clonotypes. Examining inter-genus connectivity, *Erysipelotrichaceae* (family), *Collinsella* and *Coprobacter* emerged as the most interconnected genera, sharing convergent public TCRs with multiple other genera. Many of these genera are known or suspected to have immunomodulatory effects in the gut^29–33^. Louvain community detection revealed not only independent communities containing a single genus and its associated convergent TCRs, but also other communities spanning multiple genera. Network exploration thus revealed dense core regions with overlapping associations, consistent with shared TCR binding. Additionally, we showed that MAIT α-chain clonotypes, identified by canonical V/J usage, were largely absent, consistent with their broad antigen recognition leading to lack of genus-specific signatures to be captured by this framework^34^.

Convergent TCRs were clustered by sequence similarity and we identified genus-specific TCR motifs shared across individuals. Genera such as *Porphyromonas* and *Peptoniphilus* displayed tightly clustered, high-convergence TCRs, indicative of conserved antigen recognition, whereas *Lachnospiraceae* and *Dorea* showed weaker and more diffuse clustering. Collectively,

AIRRWAS mapped TCR–microbiome interactions at both the clonotype and cluster level, identifying bacterial genera associated with convergent and shared TCR motifs. These findings suggest that specific microbial taxa may actively shape T cell repertoires. We therefore tested whether the observed associations were reproducible across independent cohorts.

### AIRRWAS identifies robust, reproducible genus-specific TCR clusters across different cohorts

To validate and prioritize TCR-bacterial genus associations identified in the AIRRWAS discovery cohort, we analyzed two independent datasets with paired gut TCR and microbiome profiles (Pu et al., CRC, n=21; Brand et al., IBD, n=16). These datasets differed in TCR source (blood vs. tissue) and microbiome profiling (16S rRNA vs. metagenomics), consistent with previous findings that microbiome-reactive TCRs can be found both in mucosal tissue and peripheral blood^14–16^. A new TCR-bacterial genus interaction network was constructed independently for each dataset using the AIRRWAS method. To examine broader antigen-driven responses, we then clustered the convergent TCRs from all cohorts together and identified motif-level convergent clusters. Clusters containing TCRs from all three datasets significantly associated to the same genus were defined as shared clusters, reflecting conserved TCR motifs (Figure 2a, Extended Data Fig. 1, Extended Data Table 1).

**Figure 2.**
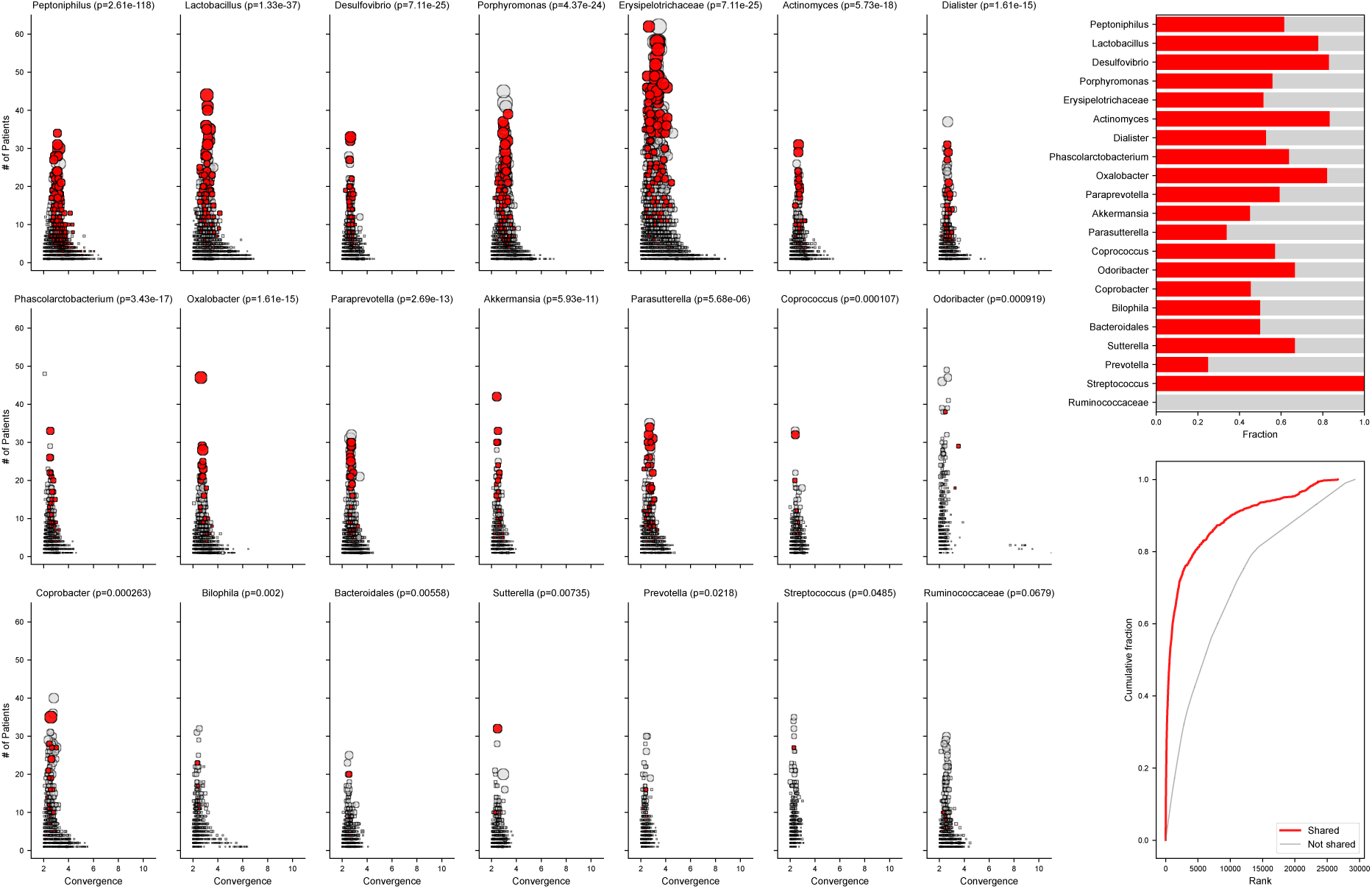
Shared convergent TCRs are enriched among highly convergent and broadly observed genus-associated clusters. **a**, Scatterplots of cluster convergence versus patient prevalence for 21 bacterial genera that yielded shared convergent clusters. Each point represents a TCR cluster, sized by the number of unique TCRs within the cluster. Shared clusters observed across all three datasets (or two for *Peptoniphilus*, absent from the Brand et al. cohort) are colored red; non-shared clusters are light grey. Red shared clusters are enriched in the top-right region, indicating high convergence and broad patient representation. The adjusted p-values from Mann-Whitney U tests comparing ranks of shared versus non-shared clusters are shown in the titles. Of the 21 genera with at least one shared cluster, 20 show statistically significant enrichment after multiple testing correction, the remaining genus approaches significance (adjusted p = 0.07). **b**, Distribution of shared clusters among the top 10% ranked genus-specific clusters across the 3 cohorts. Bars are colored according to shared (red) or non-shared (light grey) clusters. Shared clusters dominate the top for most genera, reflecting their reproducibility and prominence across datasets. **c**, Cumulative distribution of ranks across all clusters and genera. The empirical cumulative distribution function shows that shared clusters are concentrated among higher-ranked clusters relative to non-shared clusters, highlighting consistent enrichment of reproducible TCR-genus interactions.

Despite biological and technical heterogeneity, we identified 1,268 shared TCR clusters, containing 26,877 unique TCR sequences significantly convergent across 21 genera in all 3 cohorts (Extended Data Table 3). These genera, linked to convergent TCRs across datasets, include core commensals dominating healthy gut communities (*Ruminococcaceae*, *Prevotella*), classic probiotics (*Lactobacillus*), and less abundant taxa with potential immunomodulatory or pro-/anti-inflammatory roles (*Akkermansia*, *Desulfovibrio*, *Bilophila*) or opportunistic pathogens (*Porphyromonas*, *Peptoniphilus*, *Actinomyces*). This demonstrates cross-cohort reproducibility at the cluster level, meaning exact TCR sequences did not need to be shared across all datasets. The number of shared clusters per genus did not scale with total cluster counts, suggesting that some genera elicit broad, general T cell responses, whereas others drive narrower, more specific repertoires.

Ranking clusters by normalized convergence and publicity revealed that shared clusters were significantly enriched among the highest-ranked clusters for nearly all genera (20/21, Mann-Whitney U test, Benjamini-Hochberg corrected p < 0.05; Figure 2b, Extended Data Fig. 3). The remaining 15 genera did not yield any shared clusters, likely linked to their high genus prevalence. Analysis of the top 10% ranked clusters per genus (Figure 2b) and cumulative distribution of cluster ranks (Figure 2c) further confirmed that shared clusters occupy higher ranks, indicating that stronger convergence scores relate to biological relevance. These findings indicate that AIRRWAS reliably identifies both exact-matching TCRs and conserved sequence motifs across independent cohorts and sequencing techniques, reflecting biologically relevant TCR-microbiome interactions and providing a prioritized set of genus-specific TCRs for downstream functional validation.

### Convergent TCRs reveal a heterogenous microbiome-driven immune response

We next explored the distribution of T cell subtypes within convergent TCRs to reveal potential cell type-specific associations. To test this, we compared the fractional distribution of T cell subtypes between convergent and baseline TCRs and calculated log₂ fold-change (logFC) enrichment for each cohort separately (Extended Data Fig. 4). In the discovery cohort, no consistent enrichment for either CD4⁺ or CD8⁺ T cells was observed in convergent TCRs. Similarly, in the Pu et al. and Brand et al. cohorts, cell type distributions were generally balanced, without any systematic skewing towards specific cell subtypes. Instead, enrichment patterns were heterogenous and varied significantly per genus. For example, in Pu et al., most but not all genera displayed convergence biased towards CD8⁺ effector- and IEL cells, often accompanied by enrichment of CD4⁺ Th17 and/or Th1 cells, whereas CD4⁺ Treg-associated TCRs were generally depleted. Interestingly, the CD4⁺ Treg-IL17 subset^35, 36^ showed different behavior, with both enrichment and depletion depending on genus. In contrast, the Treg subset was not consistently depleted in the Brand et al. cohort, while some genera (including *Lactobacillus*, *Parasutterella*, *Erysipelotrichaceae* and *Akkermansia*) did show a depletion of non–gut-homing T cells.

Overall, no single T cell subset consistently drove genus-specific convergence across cohorts. Instead, cell type distribution within convergent TCRs was highly variable and largely genus- and cohort-specific, with genus prevalence having only minor influence. These results suggest that multiple T cell lineages contribute to genus-specific immune responses, highlighting the heterogeneity of microbiome-driven T cell recognition.

### *In vitro* bacterial stimulation validates AIRRWAS-predicted TCR-microbiome associations

To directly validate if AIRRWAS predictions are in part capturing direct T cell microbiome interactions, we performed *in vitro* stimulation of PBMCs from the discovery cohort with lysates from two bacterial species representing contrasting network profiles: *Prevotella bivia* (PB) and *Peptoniphilus lacrimalis* (PL). Stimulations were carried out for 1 day and 1 week, after which activated T cells were isolated and TCRs were sequenced (Extended Data Fig. 1, Extended Data Fig. 5a). Across all experiments, we identified 11,050 unique TCRs in PB-stimulated samples, 15,934 in PL-stimulated samples, and 621 TCRs shared between both conditions. Of these shared TCRs, 13 α-chain clonotypes (114 clones in total) were identified as MAIT cells based on V- and J-gene usage. Considerable overlap was observed between stimulated repertoires and the original discovery dataset, particularly for α chains, with matches derived from the broader cohort, patient-matched individuals and the list of convergent TCRs (Figure 3a, Extended Table 4).

**Figure 3.**
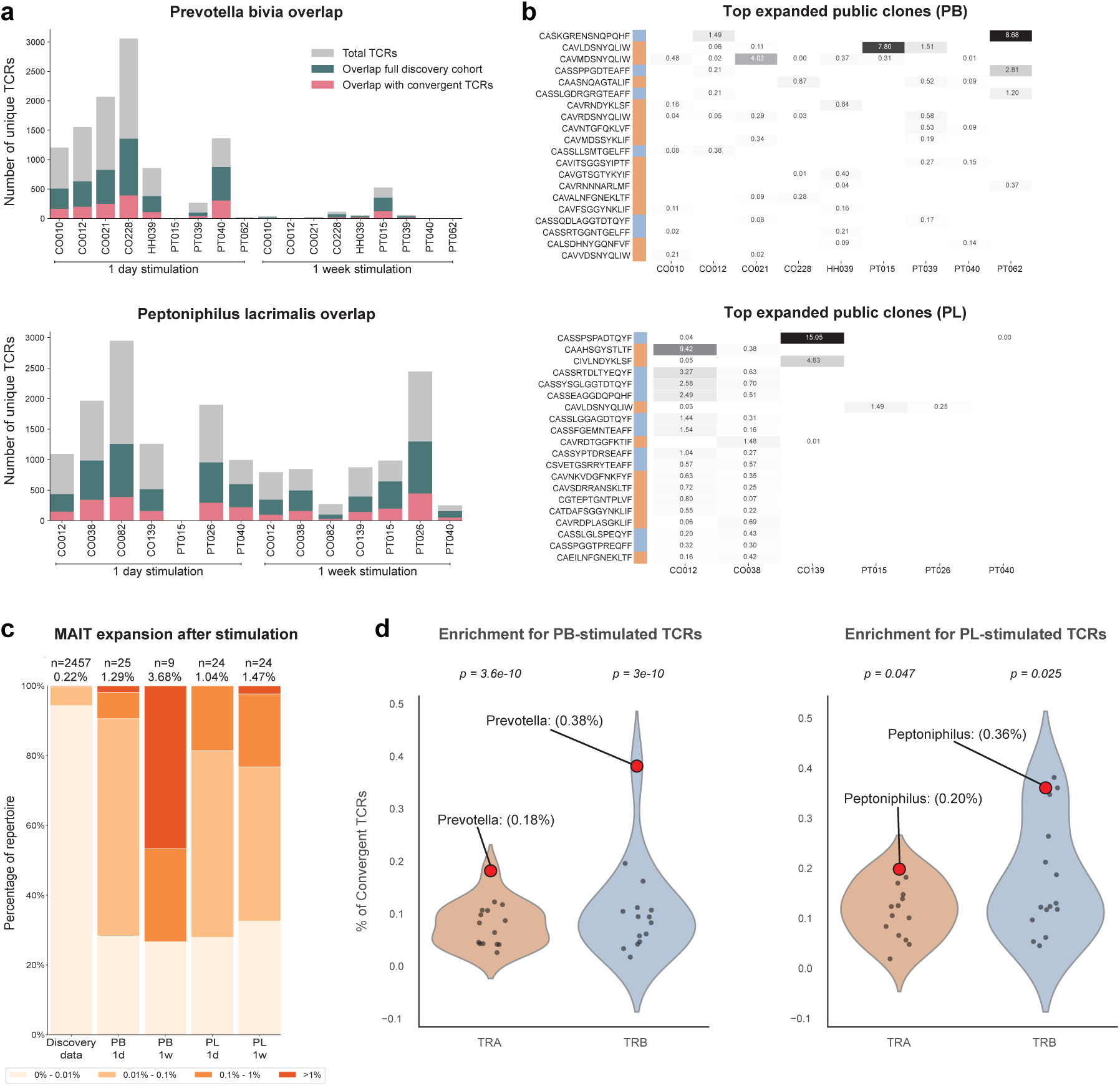
**Bacterial-stimulated TCRs were successfully predicted using AIRRWAS**. **a**, Per-patient TCR counts and overlap with the discovery cohort. Bars and colors indicate the total number of unique TCR clones detected per patient (grey), the number of TCRs that overlap with the full discovery repertoire (green) and the overlap with the set of convergent TCRs (pink). **b**, Heatmaps of relative clone frequency per sample for the top 20 most expanded public clones. Color intensity corresponds to clone abundance within the sample repertoire, highlighting recurrent clonotypes shared across multiple individuals. **c**, MAIT cell expansions in discovery and stimulated datasets. Stacked bars represent the fraction of MAIT cells within clonal expansion ranges in the discovery cohort, PB-stimulated samples or PL-stimulated samples after 1 day or 1 week stimulation. Values on top of the bars indicate the number of unique MAIT sequences and percentages indicate their contribution to the full repertoire. MAIT expansion is increased in stimulated populations as expected. **d**, Convergent TCR overlap analysis for PB- and PL-stimulated repertoires. Violin plots show relative overlap (% of genus-specific convergent TCRs) between stimulated repertoires and predicted convergent TCRs for the different genera (minimum 3000 convergent TCRs per genus). Each dot represents overlap between the stimulated TCRs and genus-specific convergent TCRs for all genera with shared clusters. Red dots indicate overlap with the correct genus-specific TCRs, with annotations indicating overlap values. Statistical significance of correct-genus overlap compared to other-genus overlap is calculated using z scores and shown above each violin plot. Convergence analysis successfully predicts genus-specific stimulated TCRs more accurately than unrelated genera.

Stimulation induced clear clonal expansion, particularly after 1 week (Extended data Fig. 5b). Combining the activated TCRs from 1 day and 1 week stimulations, PB-stimulated samples revealed 99 public clonotypes, whereas PL-stimulated samples contained 221 public clonotypes. Interestingly, PB-specific expansions were frequently observed across multiple individuals, whereas PL-specific expansions were restricted primarily to a single individual, with low-expansion in others likely reflecting high generation probability rather than antigen-specific responses (Figure 3b, Extended Data Fig. 5c+d). MAIT cells were consistently enriched after stimulation compared to the discovery dataset (Figure 3c).

Next, we tested whether the stimulated TCRs mainly overlapped with AIRRWAS-predicted genus-specific clusters for PB or PL, rather than unrelated genera. For each genus in which we identified shared clusters, we quantified the relative overlap of stimulated TCRs with the genus-specific convergent repertoire, and compared this to overlap with non-matching genera (Figure 3d). Z-scores were calculated to evaluate whether overlap with the correct genus exceeded that of non-matching overlap. The analysis was restricted to genera with shared clusters and ≥3,000 associated convergent TCRs, but including *Prevotella*. PL-stimulated TCRs showed significant overlap and enrichment for both TCR chains (α p=0.047, β p=0.025). PB-stimulated repertoires, despite its smaller set of *Prevotella*-convergent TCRs (∼1,800), still showed significant enrichment (α = 3.6e-10, β = 3e-10) for overlap between the stimulation experiment and predicted convergent clusters. Together, these findings demonstrate that TCRs from T cells that were activated by bacterial stimulation correspond to those described by the AIRRWAS method within the genus-specific convergent repertoires. This provides direct experimental validation that AIRRWAS can capture biologically meaningful, genus-specific T cell responses, even within the immense diversity of human TCR repertoires.

To further characterize cell type specificity of microbiome-associated TCR convergence, we performed a follow-up stimulation experiment using *Peptoniphilus lacrimalis* lysate on six additional samples. After 6-hour stimulation, T cells were sorted into conventional CD4+ T cells (Tconv, CD154^+^CD137^-^) and regulatory T cells (Tregs, CD154^-^CD137^+^)^37^, allowing a higher-resolution exploration of repertoire composition and overlap with the discovery dataset. Across patients, unique Tregs were approximately five-fold less abundant than Tconv cells (Extended Data Fig. 6a). Despite lower numbers, Tregs showed greater relative clonal expansion (Extended Data Fig. 6b), consistent with more focused or selective activation within the regulatory immune response. Analysis of stimulated TCRs that overlap with the convergent TCRs from AIRRWAS revealed only modest enrichment for Tregs for some bacterial genera (Log_2_FC < 1; Extended Data Fig. 6c) and no enrichment for others. Per-sample analysis showed considerable variability, with no consistent bias towards either Tregs or Tconv cells. This further supports our earlier findings that convergent TCRs are not limited to a single T cell subset but reflect a heterogenous immune response to the microbiome. The overlap of stimulated TCRs with the *Peptoniphilus*-specific convergent TCRs versus TCRs for non-matching genera revealed a significant enrichment for *Peptoniphilus*-predicted TCRs for the β chains compared to other genera (Extended Data Fig. 6d).

## Discussion

Here we describe a high-resolution framework for linking bacterial taxa to convergent TCR repertoires, integrating computational prediction with targeted *in vitro* validation. Whereas previous approaches provide important insights by associating taxonomic differential abundances with broad T cell responses^38^, AIRRWAS works at a higher resolution, identifying clonotype- and motif-level convergence patterns reproducible across independent datasets and sequencing platforms. By applying AIRRWAS to three cohorts followed by *in vitro* T cell stimulation, we discovered robust TCR-microbiome associations spanning 21 genera, that exhibited significantly higher convergence and publicity. These genus-specific convergent associations reflect clusters of similar TCRs converging on the same bacterial genus, bypassing the need for exact TCR matching across datasets. This shows that distinct repertoires can independently converge on common microbial components and highlights the potential for conserved, population-level T cell responses to the gut microbiome^10, 24, 39^.

Our results expand on growing evidence that specific microbes shape mucosal immunity in health and disease^1, 40^. For example, *Fusobacterium nucleatum* promotes inflammation and tumorigenesis in colorectal cancer^41, 42^, while *Bacteroides fragilis* has been associated with both protective and pathogenic T cell responses in colitis and inflammatory bowel disease^43, 44^. Other taxa, including *Bacteroides vulgatus, Ruminococcus gnavus*, and *Akkermansia muciniphila*, have been linked to epithelial dysfunction, autoimmunity, and metabolic disease^45–47^. The 21 genera linked to convergent TCRs in our study include both genera connected to these previously studied taxa (*Bacteroidales*, *Ruminococcaceae*, *Akkermansia*) and genera not previously implicated in immune modulation, highlighting new potential TCR-microbiome interactions. Current studies like the ones mentioned above, however often focus on single species or global outcomes, providing an incomplete view of the mucosal immunity and TCR–microbiome interactions. AIRRWAS addresses this gap by systematically mapping genus-level TCR convergence across multiple cohorts, revealing a broader landscape of microbiota-driven T cell responses that complements and extends species-focused immunological studies.

Computational predictions were supported by targeted *in vitro* stimulation experiments. T cell stimulation with *Peptoniphilus lacrimalis* lysate yielded expanded TCRs significantly enriched for convergent sequences predicted for *Peptoniphilus* in two separate experiments. *Prevotella bivia*, despite a smaller predicted convergent repertoire, also showed significant relative enrichment for both TCR chains. The follow-up stimulation experiment incorporating T cell subset sorting demonstrated that *Peptoniphilus*-activated TCRs were distributed across both conventional CD4⁺ T cells and regulatory T cells, with no preferential enrichment for either subset. This indicates that genus-specific TCR convergence is not restricted to one functional compartment. Although Tregs were less abundant overall, they exhibited greater clonal expansion, consistent with selective activation within the regulatory compartment^48, 49^. Together with cross-cohort analyses, these results show that AIRRWAS-predicted repertoires capture biologically relevant clonotypes that are preferentially activated by genus-matched stimuli, rather than randomly distributed across microbial taxa. We also showed that it captures microbiome-associated TCRs distributed across multiple functional compartments, reflecting both effector and regulatory immune responses.

Several genera identified through AIRRWAS encompass well-known species implicated in modulating immune responses in cancer, autoimmunity or chronic inflammation. The identification of convergent, public TCRs for these taxa suggests potential use as biomarkers of microbial exposure or as tools for monitoring immune responses during microbiome-targeted therapies. A major strength of AIRRWAS is its ability to detect these associations across independent cohorts and sequencing platforms, demonstrating robustness despite heterogeneity in data type and processing. Moreover, by combining our computational predictions with experimental validation, this research provides a unique link between predicted sequence-level convergence and actual functional immune responses to gut bacteria. AIRRWAS and its findings complement mechanistic studies showing how microbial products such as short-chain fatty acids or polysaccharide A can influence T cell differentiation and immune tolerance, highlighting the multi-level regulation of mucosal immunity by commensal and pathogenic microbes^50–53^.

Despite these advances, several limitations must be considered. Genera with very high prevalence displayed fewer convergent TCRs and increased publicity among associated clonotypes, suggesting that high-publicity sequences may partly reflect generation-probability biases rather than antigen-driven selection. For these taxa, the reduced availability of negative samples may create false positive associations. In addition, stimulation experiments using whole lysates, while valuable for validation, cannot resolve the precise antigenic epitopes driving these immune responses. Pinpointing cognate peptides will require complementary approaches such as peptide–MHC multimer staining, high-throughput antigen screens and immunopeptidomics. Finally, although AIRRWAS demonstrated robustness across three heterogeneous cohorts, we expect that the constructed network could contain many false positive interactions, which can be further resolved with more (larger) or additional complementary datasets. Nevertheless, to our knowledge AIRRWAS represents the first network-based framework capable of predicting TCR–bacteria interactions at this resolution.

Taken together, our findings demonstrate that genus-specific TCR convergence is detectable, reproducible and experimentally validated across independent datasets and immune compartments. By linking bacterial taxa to TCR repertoires, AIRRWAS provides a generalizable framework for finally revealing the hidden TCR-microbiome interaction network. Shared clusters with high convergence and publicity represent promising candidates for public TCR biomarkers, while future integration with single-cell transcriptomics, structural modeling and spatial or temporal TCR repertoire tracking could connect sequence-based predictions to functional immune states. More broadly, this work lays the foundation for mechanistic and translational studies at the interface of the microbiome and adaptive immunity, with potential applications in biomarker discovery, disease monitoring, and the development of microbial or peptide-based immunotherapies.

## Online Methods

### Discovery cohort sample collection

A total of 74 patient samples were collected with written informed consent and ethics approval from the Antwerp University Hospital IRB (reference number 20/02/003), including PBMC bulk TCR sequencing and 16S rRNA stool sequencing data. These individuals, age range 15-78 (mean ± sd: 56.01 ± 12.66) were recruited and donated blood between September 2020 and July 2021. Extensive metadata was also collected, including sex (39 female, 35 male), severe acute respiratory syndrome coronavirus 2 (SARS-CoV-2) infection, Cytomegalovirus (CMV) IgG levels, Body-Mass Index (28.09 ± 5.96), Beck Depression Inventory score, Beck Anxiety Inventory score, occurrence of a previous SARS-CoV-2 infection, long Covid symptoms, 25-hydroxy Vitamin D and current smoker status. Further detailed information regarding these characteristics, additional patient data and sample processing can be found in Ha et al.^54^ and Affaticati et al.^55^. In short, whole blood samples were collected in lithium-heparin, processed using SepMate-50 tubes and finally stored in Liquid Nitrogen. CD4^+^ and CD8^+^ cells were separated using magnetic beads after which bulk TCR sequencing was performed [Qiagen TCR kit] and TCR sequences were identified using MiXCR^56^ v3.0.13 with the default input parameters.

Sequencing of the 16S rRNA V4 region in stool samples was performed to characterize the bacterial strains as described in Affaticati et al.^55^. Raw 16S rRNA files were processed using DADA2^57^ v1.8.0 and Tidytacos^58^ to identify amplicon sequence variants (ASVs), add taxonomic annotation using the EzBioCloud^59^ 2018.5 database and calculate their abundance within each sample. This was used as input for the AIRRWAS pipeline, using the microbiome frequency at the genus level.

### TCR diversity and epitope prediction

TCR sequencing data was processed to remove non-canonical CDR3 sequences, exclude non-functional gene segments and standardize sequences according to ImMunoGeneTics (IMGT) nomenclature^60^. Unique clonotypes were defined by distinct combinations of CDR3 nucleotide sequence, CDR3 amino acid sequence, V- and J-gene and cell type per patient. Duplicate TCR entries were collapsed and patient-specific clone counts and relative frequencies were aggregated per clonotype. The TCR richness and alpha diversity profiles within each repertoire were calculated, including Pielou’s Evenness, Gini-Simpson diversity, Gini coefficient and the DE25^61^ value using scikit-bio v0.5.9. Differences in diversity across T cell subgroups including TCR α (TRA) and TCR β (TRB) chains and T cell subtypes (CD4^+^ and CD8^+^), were compared. Beta diversity between repertoires was determined by calculating the Bray-Curtis dissimilarity with SciPy v1.11.3 and visualized using t-distributed stochastic neighbor embedding (t-SNE). TCR epitope specificity was predicted using the ImmuneWatch DETECT algorithm^62^ v1.4.0.

Predictions were deemed significant at the default threshold of 0.2 and the epitopes were further grouped in general categories: “Viral”, “Human”, “Microbial” and “Synthetic”.

### Taxonomic profiling of the stool samples

Microbiome taxonomic profiling was performed using DADA2 v1.8.0 to identify individual ASVs from 16S rRNA sequencing of 74 samples, which resulted in 3803 unique ASVs. Individual ASVs were aggregated at the genus level if taxonomic annotation was available. For ASVs without a genus-level annotation, aggregation was performed at the next lowest available taxonomic rank (typically family). For clarity, all resulting groups were referred to as “genera” throughout the manuscript, although some may correspond to family-level classifications. Relative abundances per genus were then calculated per sample. Differential genus-level abundances were visualized for the top nine most abundant genera, while remaining taxa were aggregated into an “Other” category. To evaluate the global structure of the microbiome across samples, Bray-Curtis dissimilarity was calculated and projected using t-SNE. Genus prevalence across the cohort was determined by calculating the proportion of samples in which each genus was present versus absent.

### TCR-microbiome association analysis

We developed and applied the novel Adaptive Immune Receptor Repertoire Association Study (AIRRWAS) to identify associations between TCR repertoires and the gut microbiome (Extended Data Fig. 1). AIRRWAS first uses TRIASSIC^63^ [v0.2.0, Python v3.10.14] to identify convergent TCRs linked to a bacterial genus. TRIASSIC quantifies convergence by counting similar clonotypes that arise independently, defined as: (i) identical clonotypes in different individuals, (ii) clonotypes with distinct nucleotide sequences encoding the same amino acid sequence regardless of individual and (iii) highly similar clonotypes (TCRdist score ≤ 12.5), regardless of the individual. For each TCR, convergence events are calculated within its 2048 most similar clonotypes, as identified by TCRdist^64^. Enrichment is determined using a two-tailed Fisher’s exact test comparing samples containing the genus versus those without.

A total of 36 different genera were analyzed using the AIRRWAS pipeline. For each genus, TCR convergence was calculated, followed by enrichment testing to extract significantly convergent clonotypes. Convergence and publicity of the individual convergent TCRs was compared across T cell subtypes (CD4^+^, CD8^+^) and chain types (α, β). Next, AIRRWAS selects statistically significant clonotypes (convergence > 2, p ≤ 0.05) and clusters them per genus by sequence similarity (Hamming distance = 1) using ClusTCR^65^ v1.0.2. For each cluster, convergence, enrichment and publicity were compared across genera to identify genus-specific or patient-enriched clusters. Based on these results, AIRRWAS generates an interactive network visualization (Pyvis v0.3.2) of TCR-microbiome associations, with nodes representing TCR clusters or bacterial genera and edges indicating statistically significant convergence. Communities, representing groups of nodes more densely connected to each other than to the rest of the network, were identified using the Louvain community algorithm (python-louvain v0.16).

### Additional cohort collection

To confirm the generalizability of our findings, we analyzed two independent cohorts with paired TCR and microbiome data, differing in sample type and sequencing approach. One cohort was selected for metagenomic sequencing, while the other contained combined tissue-derived TCR and 16S rRNA data, in contrast to the discovery cohort’s blood-derived TCR and stool 16S rRNA data. Comparative analyses across three independent cohorts identified overlapping and cohort-specific TCR-microbiome associations.

#### Brand et al. twin IBD cohort

This cohort is derived from the prospective Dutch study “Twin cohort for the study of (pre)clinical inflammatory bowel disease in the Netherlands” (TWIN-IBD; Dutch Trial Register: NL6187). For this study, TCR and microbiome data of 4 concordant and 4 discordant monozygotic twin pairs (n = 16) for Crohn’s disease were included. Peripheral blood T cells were sorted via FACS into Tregs (CD3^+^CD8^−^CD4^+^CD25^+^CD127^low^), gut-homing memory CD4^+^ T cells (CD3^+^CD8^−^CD4^+^CD25^−^CCR7^+/−^CD45RA^−^Integrin-α4β7^+^), and non-gut-homing memory CD4^+^ T cells (CD3^+^CD8^−^CD4^+^CD25^−^CCR7^+/−^CD45RA^−^Integrin-α4β7^−^). Bulk TCR sequencing was performed on sorted T cell populations as described in Brand et al*. (in preparation)*. TCR sequencing data was processed using the same pipeline as the discovery cohort to ensure consistency.

Fecal samples were collected concurrently for microbial DNA extraction and whole-genome shotgun metagenomic sequencing as described in Brand et al.^66^. in short, taxonomic profiling was conducted with MetaPhlAn2 (v2.7.2, mpa_v20_m200), excluding non-bacterial sequences and maintaining species-level annotations. Species were aggregated at the genus level and paired TCR and metagenomic data were then processed through AIRRWAS, substituting 16S rRNA data with metagenomic profiles.

#### Pu et al. single cell CRC cohort

The Pu et al. cohort comprised 21 CRC patients with paired tissue TCR single-cell sequencing and tissue 16S rRNA sequencing from tumors and adjacent non-malignant tissue. Details on patient recruitment and sample processing are described in Pu et al., (*in preparation*). Tumor and adjacent tissues were collected via curative surgical resection, fresh-frozen and homogenized for DNA extraction. Single-cell RNA sequencing of T cells was performed on the 10x Genomics Chromium platform, and T cell subtypes were annotated with STEGO.R^67^. Microbiome composition was analyzed through 16S rRNA sequencing and taxonomic assignment using DADA2, after which abundances were collapsed at the genus level. Tumor microsatellite stability and immune consensus molecular subtypes were determined. Tissue-specific TCR-microbiome interaction were processed, parsed and interaction networks were constructed using AIRRWAS.

### Cross-cohort matching of convergent TCRs and cell types

AIRRWAS was also applied to the additional independent cohorts for the same 36 bacterial genera. For each cohort, TCR convergence was calculated independently using TRIASSIC, with convergent TCRs defined as clonotypes with convergence > 2 and p ≤ 0.052 for a given genus. To identify convergent TCR-genus patterns that consistently occurred across cohorts, we used two complementary approaches: (1) exact sequence matching to identify TCR clonotypes convergent for the same genus across all datasets, and (2) motif-level matching, where all convergent TCRs were combined and clustered per genus using ClusTCR. Clusters containing TCRs from all three datasets were defined as shared clusters, irrespective of cluster size. For each cluster, mean convergence and the number of unique patients per cluster were calculated. To evaluate whether shared clusters represented stronger biological signals (i.e. higher convergence and publicity), clusters were ranked within each genus according to the sum of their normalized mean convergence and publicity. Shared clusters were then compared to non-shared clusters using a Mann-Whitney U test, with Benjamini-Hochberg multiple testing correction for genera containing at least one shared cluster. To further visualize the distribution of ranks, empirical cumulative distribution functions (ECDFs) were generated for shared versus non-shared clusters.

To explore the cellular origin of convergent TCRs, we quantified the fraction of each T cell subtype within each genus that yielded shared clusters. For each dataset, the cell type of convergent TCRs was compared to the baseline set of TCRs. Absolute counts per cell subtype were normalized by the total number of cells for that T cell subtype in the baseline repertoire. For each genus, fold changes in convergent versus baseline counts were calculated and statistical significance was tested using Fisher’s exact test (for two cell types) or a chi-squared test (for more than two cell types). Resulting fold changes were visualized as a heatmap, combining all three datasets, with hierarchical clustering applied to the genera to group together similar patterns.

### T cell stimulation with *Prevotella bivia* and *Peptoniphilus lacrimalis*

To validate predicted TCR-microbiome associations, we stimulated patient-derived T cells with lysates from *Prevotella bivia* (PB) and *Peptoniphilus lacrimalis* (PL) as an example. These bacteria represent species from genera with convergent TCRs and contrasting network patterns. 100 µl *Prevotella Bivia* (BEI Resources, Strain GED7880) and *Peptoniphilus Lacrimalis* (BEI Resources, strain DNF00528) were cultivated on tryptic soy agar (TSA) with 5% defibrinated sheep blood and incubated for 2 days (PB) and 5 days (PL) at 37°C in anaerobic conditions. All handlings were performed in a hypoxystation. Finally, bacteria were gently scraped from the agar and suspended in 10 mL cold PBS. The CFU was calculated using a dilution series on TSA with 5% defibrinated sheep blood. Centrifugation at 800 x g for 25 min at 4°C for PB and 2000 x g for 25 min at 4°C for PL was done, supernatant was removed and pellets dissolved in 1 mL cold sterile water. For the preparation of bacterial lysates, bacteria were frozen in liquid nitrogen for 5 minutes, defrosted by incubation for 10 minutes at 37°C and then sonicated three times with 10 minute recovery periods on ice in between sonication steps. Bacterial fractions were again snap-frozen in liquid nitrogen for 5 min, defrosted by incubation for 10 minutes at 37°C and centrifuged at 3500 x g for 20 min at 4°C. Supernatant was collected and passed through a 0.22 µm filter and stored at −80°C in aliquots. PBMCs from 13 donors in the discovery cohort were stimulated with these lysates in two different culture protocols: for 24 hours and 7 days. Six samples were stimulated with PB lysate, four with PL lysate and three were stimulated with both lysates separately. Activated CD154^high^OX40^high^CD4^+^ and CD137^high^CD69^high^CD8^+^ T cells were isolated using FACS (FACSAria II, BD Biosciences) and subjected to bulk TCR sequencing using the QIAseq Immune Repertoire RNA Library Kit (Qiagen). TCR repertoires post-stimulation were compared to network-predicted convergent TCRs for the corresponding genera. Public TCRs were quantified, clonal expansion was determined and stimulation-related clusters were identified using ClusTCR. Exact sequence overlap (i.e., the full CDR3 amino acid sequence is identical) between PB- or PL-stimulated TCRs and the original discovery cohort was calculated separately for α and β chains. Overlap was determined at three levels, (1) overlap with the full discovery cohort, (2) patient-matched overlap within the correct patient of origin and (3) overlap with only the set of significant convergent TCRs across all genera. This last genus-specific overlap was also calculated as the percentage of overlapping TCRs normalized by the total number of convergent TCRs identified for that genus. These values were compared with overlap to convergent TCRs from all other genera with shared clusters. Enrichment significance was assessed using z-scores to confirm whether stimulation preferentially activated and expanded network-predicted TCRs for the corresponding genus, with lower activation for other genera.

### PL T cell stimulation and T cell subtype sorting

To assess T cell subset-specific responses among convergent TCRs, we stimulated samples again with PL and sorted CD4⁺ T cells into conventional (Tconv; CD154^high^CD137^low^CD4^+^) and regulatory T cells (Tregs; CD154^low^CD137^high^CD4^+^) as described elsewhere^37^. A subset of PBMC samples (n = 6) from the discovery cohort was used. PBMCs were stimulated with PL lysate prepared as described above and activated Tconv cells and Treg cells were isolated using FACS (Aurora CS, Cytek Biosciences) after 6 hours of stimulation. Activated TCRs were subjected to bulk TCR sequencing using the Throughput-Intensive Rapid TCR Library sequencing protocol (TIRTL-seq)^68^.

PL-stimulated TCR repertoires from each T cell subset were compared to convergent TCRs for *Peptoniphilus* identified in the discovery cohort. We quantified differences in clonotype abundance, clonal expansion, cell type preference among the convergent TCRs and overlap with the same and all other genera in the network (as described above), providing subset-resolved validation of TCR-microbiome associations.

## Acknowledgments

We would like to thank Karin Peeters, Joke Vereecken, Maria Matteijssens, Jolien Schippers, Hans de Reu and Sam Van Goethem for their help in participant recruitment and sampling. We like to thank Nele Van de Vliet, Ines Tuyaerts and Sam Baekelants for their help with 16S microbiome sequencing.

We would also like to acknowledge BEI Resources as the following reagents were obtained through BEI Resources, NIAID, NIH as part of the Human Microbiome Project: (1) *Prevotella bivia*, Strain GED7880, HM-1270. (3) *Peptoniphilus lacrimalis*, Strain DNF00528, HM-1161.

## Funding

The following founding sources are acknowledged:

- Research Foundation Flanders (FWO)

- 1SH6624N (V.V.D), 1SH3924N (F.A.), 1861219N (B.Og.), G0H4520N (B.Og., K.L., P.M.), G031222N (T.G., S.L.), G0C9620N (S.V., S.T.)

- European Union’s Horizon 2020 research and innovation programme grant agreement 851752-CELLULO-EPI (B.Og.) & 852600-Lacto-Be (S.L.)

- Flemish Interuniversity Council iBOF MIMICRY grant (K. L., S.V., S.T.)

- Chan Zuckerberg Initiative (P. M., K. L., B. Og.).

- The Netherlands Organisation for Health Research and Development (ZonMw), VICI grant 09150182010023 (F.v.W.)

- Alexandre Suerman program for MD and PhD candidates of the University Medical Center Utrecht, the Netherlands. (E.C.B.)

- BOF-FKO mandate FKO/21/002 (S.T.)

## Author contributions

Conceptualization: R.V., P.M., B.Og.

Methodology: R.V., V.V.D., I.D.B., T.G., S.L. F.A., N.D.R., P.D.

Investigation: R.V.

Funding acquisition: P.M., B.Og., S.L.

Validation: M.K.H., H.D.R., E.B., M.K., N.D.R, P.D., B.Og.

Visualization: R.V.

Project administration: P.M., B.Og.,

Resources: N.D.R, P.D., T.P., E.C.B., B. Ol., F.v.W., S.V., S.T, S..L.

Supervision: P.M., B.Og. Writing—original draft: R.V., P.M., B.Og. Writing—review and editing: all authors.

## Competing interests

E.C.B. is co-applicant on an Investigator Initiated research grant of Pfizer. S.L. is an academic board member of ISAPP (International Scientific Association on Probiotic and Prebiotics), has received research funding from probiotic companies and has served as consultant for Freya Biosciences, YUN and Yakult, but unrelated to this research.

B. Ol. Has received grants from Takeda, Alpha Sigma, Pfizer and Abbvie. He is also a member of advisory boards at Abbvie, Janssen, Pfizer, Ferring, BMS, Lilly, Takeda and Galapagos. F.v.W. has previously served as a speaker and/or consultant for Janssen, Johnson & Johnson, and Takeda. She has received research funding from LEO Pharma, Takeda, Galapagos, Regeneron, Sanofi, and Bristol-Myers Squibb, all unrelated to this research. P.D has received research funding from and/or has ongoing contracts with MSD, Pfizer, Gilead, and GSK, but unrelated to this research. P.M., K.L., and B.Og. are shareholders and board members of ImmuneWatch BV.

## Data and materials availability

All data supporting the findings in the paper are available within the main text and/or the Extended Materials. The sequencing data from the discovery cohort have been deposited at https://github.com/fabio-affaticati/activ_covid-tcell-omics. Experimental data generated during the validation experiments in this paper will be deposited on zenodo and made publicly available upon acceptance of the manuscript.

Data from the independent cohorts used to validate our findings are available as follows: (1) IBD twins study: The microbiome metagenomics data is available in *Brand et al.* (DOI: 10.1053/j.gastro.2021.01.030) and the TCR sequencing data is available in *Brand et al.* (in preparation). (2) Colorectal cancer single cell dataset: The microbiome data is not publicly available at this moment. The TCR sequencing data is available in *Pu et al.* (in preparation).

All code and analysis scripts developed for this study will be made publicly available upon acceptance of the manuscript at https://github.com/RomiVandoren/AIRRWAS.

## Supplementary figures

**Extended Data Fig. 1.**
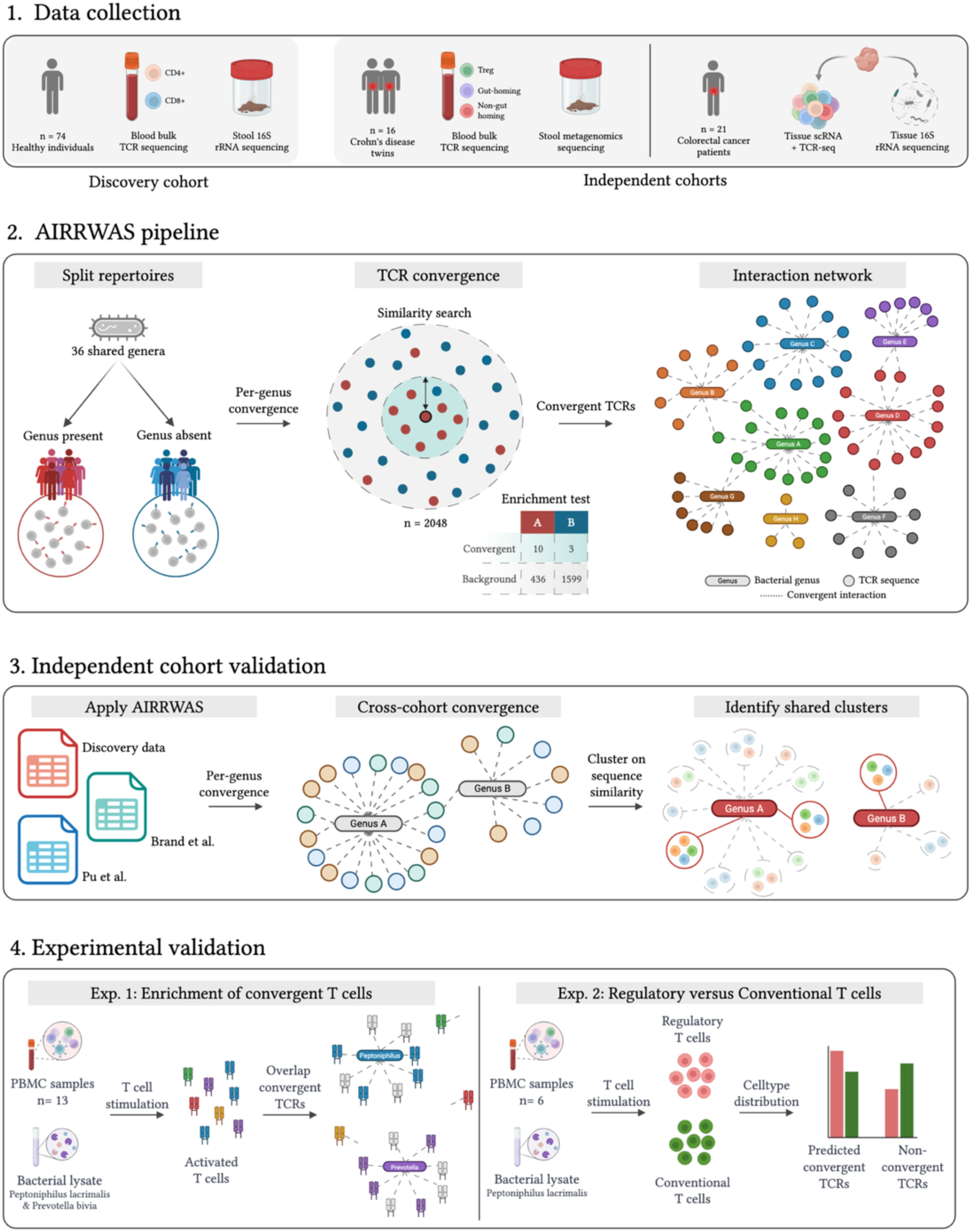
Overview of the AIRRWAS framework for generating and validating the TCR-microbiome interaction network. (1) Overview of the sample numbers and data types from the discovery cohort and two independent cohorts. **(2)** The AIRRWAS pipeline, first split the TCR data into positive and negative groups per-genus, next calculate TCR convergence using TRIASSIC and generate TCR-microbiome interaction network with convergent TCRs. **(3)** The AIRRWAS pipeline was applied to additional independent datasets, testing per-genus convergence across cohorts. Clustering of convergent TCRs identified shared sequence motifs linked to the same bacterial genus across independent datasets. **(4)** *In vitro* T cell stimulation assays to confirm computational predictions. Exp. 1: Stimulation with *Peptoniphilus lacrimalis* and *Prevotella bivia* lysates to identify enrichment for AIRRWAS-predicted convergent TCRs. Exp. 2: Stimulation with *Peptoniphilus lacrimalis* followed by subset sorting to explore distribution of *Peptoniphilus*-associated TCRs across regulatory and conventional CD4⁺ T cells.

**Extended Data Fig. 2.**
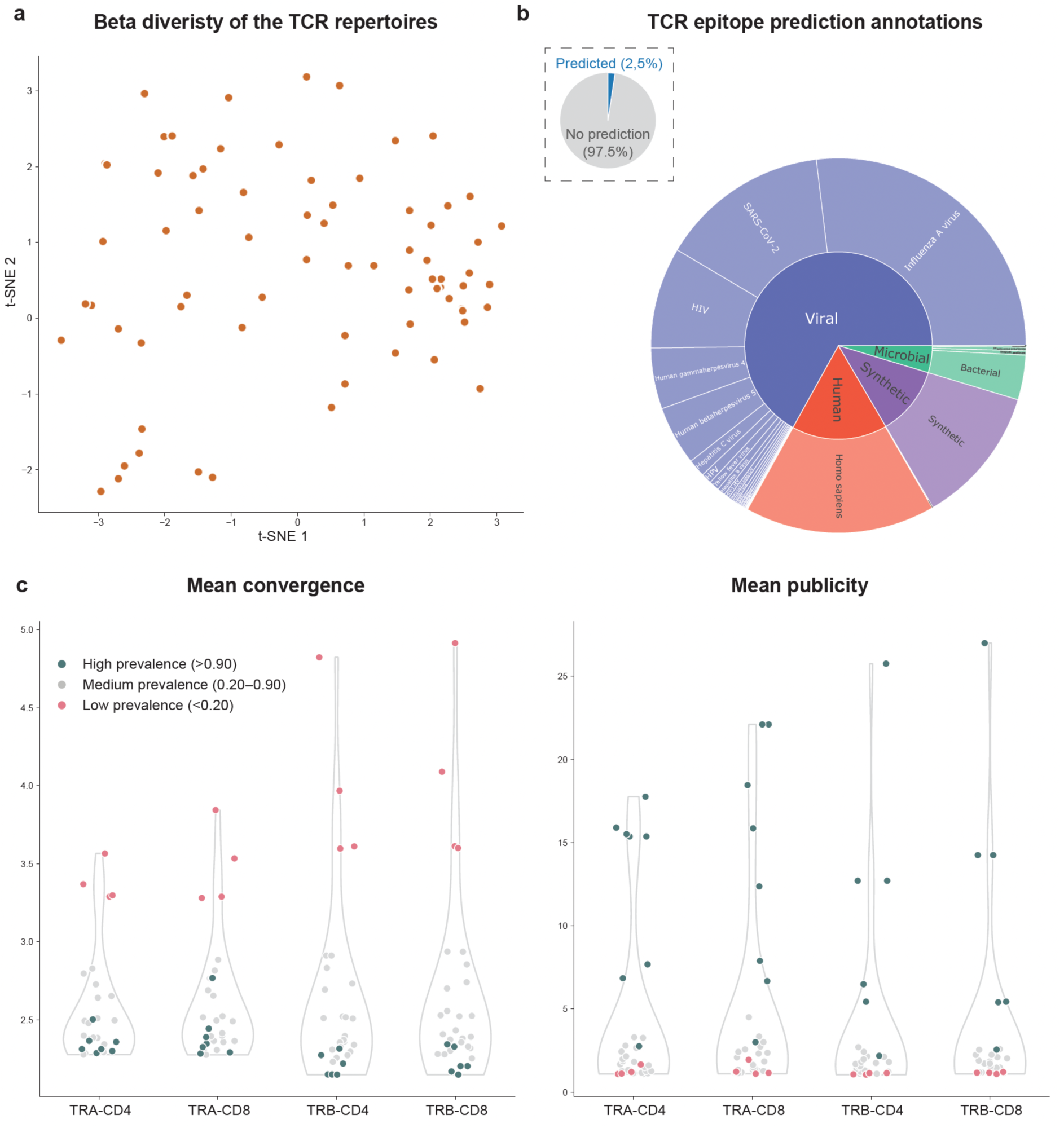
Overview of TCR repertoire diversity, epitope predictions and convergence metrics. **a**, t-SNE of TCR repertoire beta diversity (Bray-Curtis) for all discovery cohort samples. Each dot represents a sample and dots are evenly distributed across the space. **b,** Predicted TCR epitope specificity by major epitope grouping and epitope species. The inset pie shows the percentage of TCRs with significant epitope predictions (2.5%) versus those without prediction (97.5%). The inner ring shows main epitope groups (Viral, Human, Microbial, Synthetic), while the outer ring represents epitope species. **c**, Mean convergence and mean publicity of the convergent TCRs per genus (each dot is a different genus). High prevalent genera (green) show a lower mean convergence and higher publicity. Low convergent genera (pink) show a higher mean convergence and lower publicity of the response. TRA = TCR α chain, TRB = TCR β chain

**Extended Data Fig. 3.**
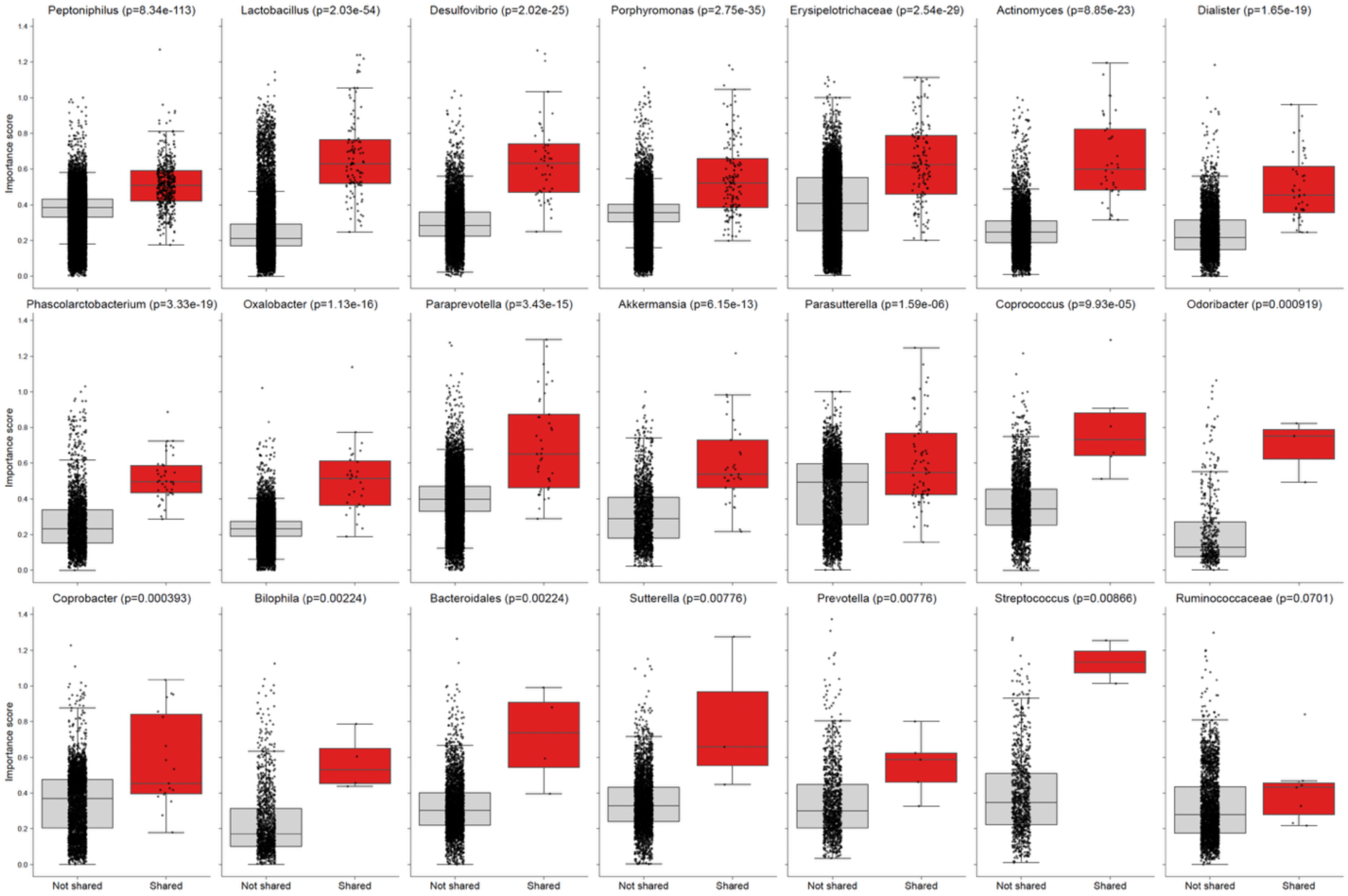
Sum rank enrichment of shared clusters for convergence and publicity. Boxplots of the importance score (used for ranking) for clusters across all genera with shared clusters. The importance score is the sum of mean convergence and publicity per cluster. Shared clusters (red) are significantly more enriched for high importance scores compared to non-shared clusters (light grey), indicating that shared clusters tend to have higher convergence and are present in more patients. All p-values were calculated using a one-sided Mann-Whitney U test and remain significant after FDR correction, except for the last genus (p = 0.07), which shows a trend toward significance.

**Extended Data Fig. 4.**
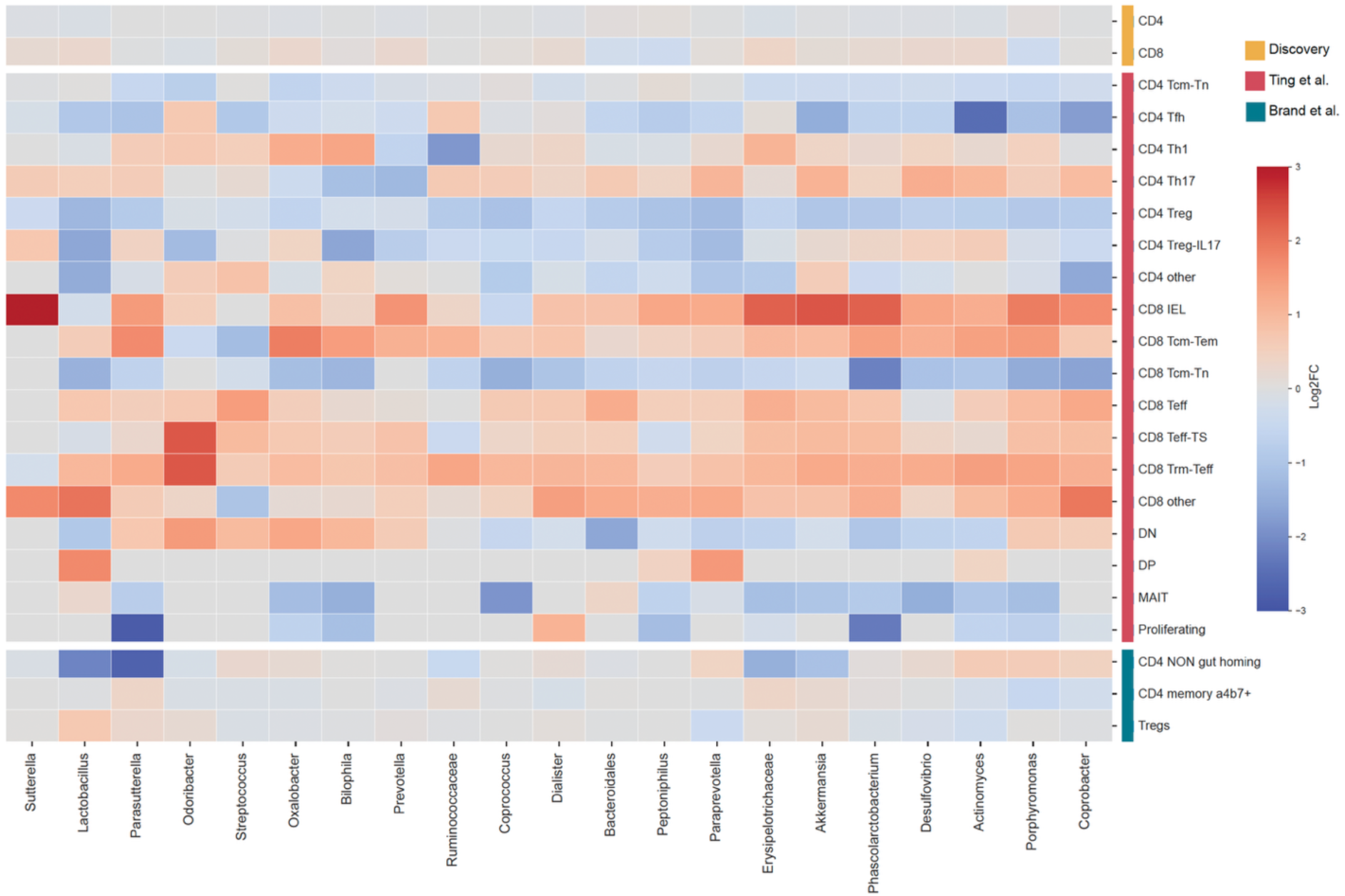
T cell subtype preferences in convergent TCRs. Heatmap of log₂ fold-change enrichment of T cell subtypes within convergent TCRs relative to the baseline repertoire across three cohorts for the 21 genera that had associated shared TCR clusters. Rows represent T cell subtypes grouped by dataset and columns represent microbial genera. In the discovery cohort (yellow), convergent TCRs were not consistently enriched for either CD4⁺ or CD8⁺ T cell subtypes. In the Pu et al. cohort (red), single cells were annotated with T cell subtypes using STEGO.R. Genus-specific patterns could be seen, but these would not transfer across all genera. This included CD8⁺ effector and IEL enrichment with concurrent CD4⁺ Th17/Th1 bias for most but not all genera. In addition, MAIT cells and CD4⁺ Tregs were generally depleted, however, Treg-IL17 had variable enrichment. The Brand et al. cohort (blue) showed no clear depletion of Tregs but non–gut-homing CD4⁺ T cells for genera such as *Lactobacillus, Parasutterella, Erysipelotrichaceae* and *Akkermansia* did show depletion, without strong enrichment of other subsets. Across all datasets, no single T cell subtype was consistently linked to the identified convergent TCRs, indicating that genus-specific TCR convergence arises from diverse and context-dependent T cell subtypes.

**Extended Data Fig. 5.**
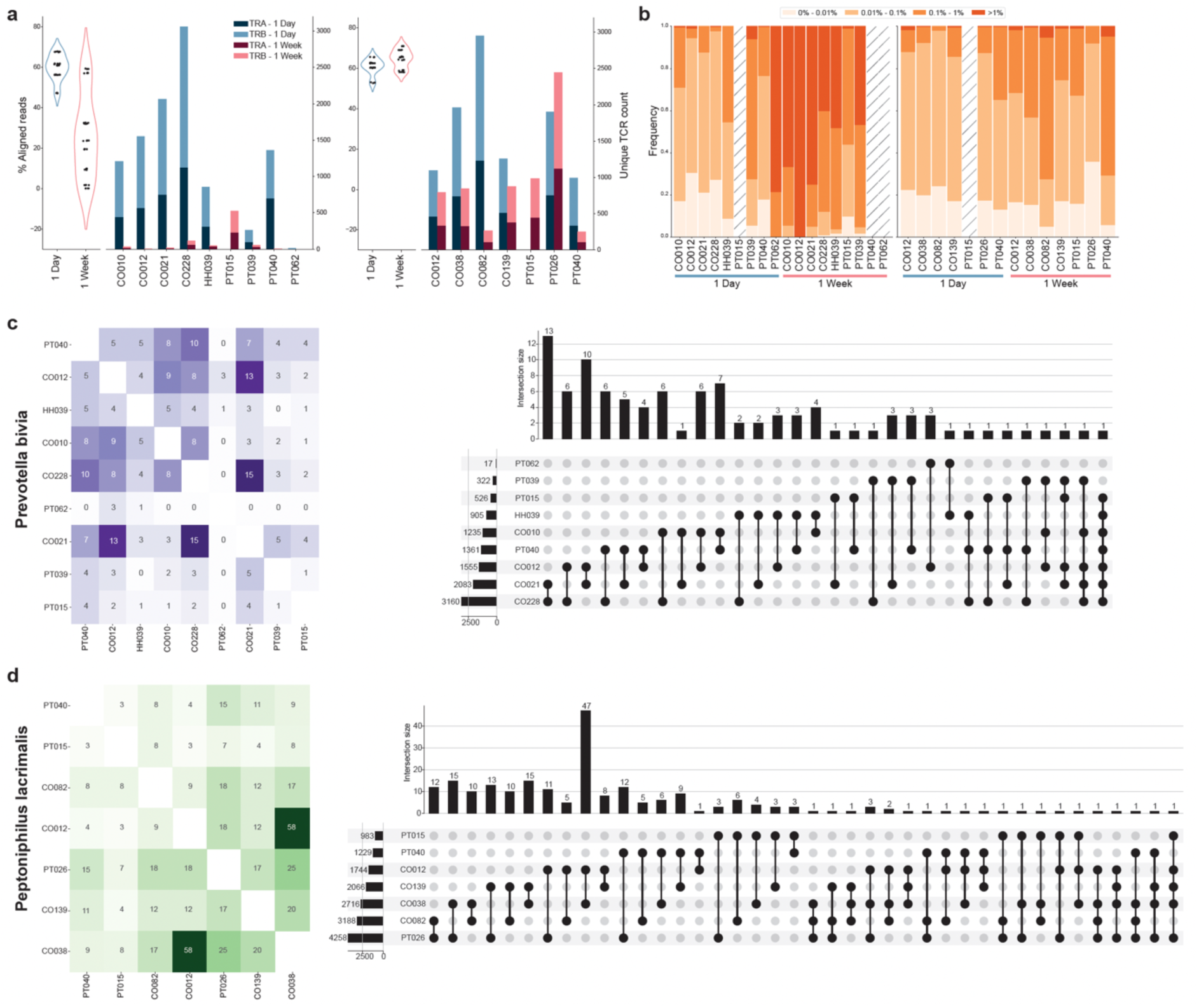
Stimulation of PBMCs with *Prevotella bivia* and *Peptoniphilus lacrimalis* lysate. **a**, Violin plot of the percentage of successfully aligned reads per sample. This shows that PB 1-week samples have limited aligned reads. Bar plots display the number of unique TCRs per patient and condition, separated by α (TRA) and β (TRB) chains. **b**, Stacked bar plots of relative clonal expansion per sample and condition for PB and PL. We observe a moderately higher clonal expansion at 1 week compared to 1 day. This is most obvious in PB 1 week, which is likely influenced by the very limited repertoire sizes. **c**, Heatmap showing absolute numbers of overlapping TCRs across the combined 1 day and 1 week repertoires of patients stimulated with PB. Numbers indicate absolute overlap, the accompanying UpSet plot summarizes public TCRs shared by two or more samples, with the most public TCR being shared between 7 different individuals. **d**, Equivalent heatmap and UpSet plot for PL, showing broader TCR sharing and more public clones shared across patients. TRA = TCR α chain, TRB = TCR β chain, PB = *Prevotella bivia*, PL = *Peptoniphilus lacrimalis*

**Extended Data Fig. 6.**
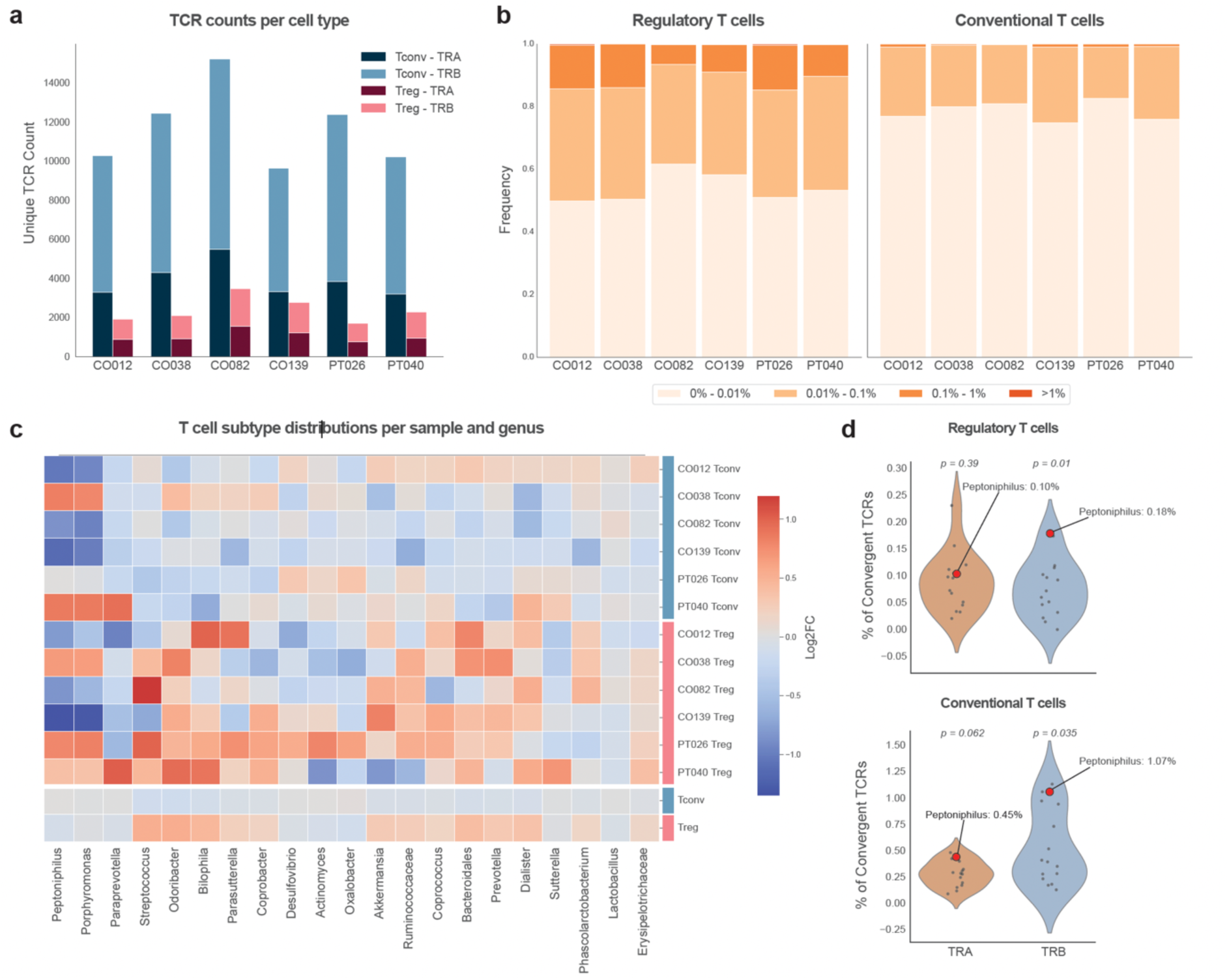
Predicted convergent TCRs do not show bias towards conventional or regulatory T cells. **a**, Bar plots of unique TCR sequences after *Peptoniphilus lacrimalis* (PL) stimulation, stratified by TCR chain (TRA, TRB) and T-cell subtype (Treg, Tconv). Tregs were approximately five-fold less abundant than Tconv. **b**, Relative clonal expansion by T cell subtype, demonstrating proportionally greater expansion among Tregs despite their lower number of unique TCRs. **c**, Heatmap of log₂ fold change between TCRs overlapping with convergent TCRs and those that don’t overlap, for Treg versus Tconv repertoires, shown per genus and per individual sample, as well as aggregated across samples. Patterns vary per genus and sample, with no consistent enrichment for either subset among stimulated TCRs overlapping with the convergent TCRs from the discovery cohort. **d**, Violin plots (Tregs, top; Tconv, bottom) for TRA and TRB chains, showing the degree of overlap with convergent TCRs predicted for PL (red) versus all other genera with shared clusters (grey). Significant enrichment is observed for TRB chains within PL-specific convergent TCRs relative to other genera, indicating that AIRRWAS accurately prioritizes PL-specific TCRs beyond random chance. TRA, TCR α chain; TRB, TCR β chain; Tconv, conventional T cells; Treg, regulatory T cells.

